# Heuristic Multi-site Optimization for Protein Sequence Design using Masked Protein Language Models

**DOI:** 10.1101/2025.07.31.668012

**Authors:** Lijuan Wang, Yuze Wang, Chen Qiu, Liwei Xiao, Xianliang Liu, Junjie Chen

## Abstract

Protein sequence design for tailored functional properties is a fundamental task in protein engineering, with critical applications in drug discovery and therapeutic development. Efficient navigation of the combinatorial vastness of protein sequence space to identify functional variants remains a formidable challenge. Conventional approaches, which predominantly rely on template-based local search or single-residue mutagenesis, are constrained by their susceptibility to local optima and their potential risk of destabilizing native structural stability. In this study, we introduce ProtHMSO, a heuristic multi-site optimization framework leveraging masked protein language models (ProtLMs) for context-aware sequence exploration. ProtHMSO mimics natural evolutionary mechanisms by employing ProtLM-derived substitution probabilities to guide heuristic searches for synergistic mutations, thereby constraining combinatorial search spaces through evolutionary and biophysical priors. Given a starting sequence and a predictive fitness model, ProtHMSO heuristically generates high-fitness multi-site variants under evolutionary constraints. Besides, ProtHMSO can also be as a modular plugin that enhances the convergence efficiency of genetic algorithms (GAs) and Monte Carlo tree search (MCTS) by integrating evolutionary priors into exploration strategies. Benchmark experiments demonstrate that protein sequences generated by ProtHMSO exhibit superior functional performance and closer alignment with natural sequence distribution, compared with state-of-the-art methods. These advancements highlight that ProtHMSO has strong potential and compatibility to accelerate functional protein discovery, offering a robust framework for efficient and context-aware exploration of protein sequence space.

**Author summary:** To address the challenge of efficiently discovering functional new proteins in protein engineering due to the vast sequence space, and to overcome the limitations of traditional evolutionary algorithms that rely on blind random mutagenesis, resulting in inefficiency and prone to structural destabilization, we proposed a heuristic multi-site optimization framework, ProtHMSO. Its core concept is to leverage the powerful contextual prediction capabilities of masked protein language models (such as ESM-2) to guide sequence mutagenesis. By predicting amino acid substitutions at specific sites that are consistent with evolutionary laws and biophysical priors, ProtHMSO narrows the exploration scope from the vast combinatorial space to a small number of high-potential candidate sequences, achieving intelligent and efficient optimization of protein sequences. Furthermore, ProtHMSO is not just a standalone algorithm, but also a plug-and-play “enhancement module”. By embedding it into a genetic algorithm (GA) and a Monte Carlo tree search (MCTS), it replaces the random mutation operator in the former with its intelligent mutation and guides the tree expansion process in the latter. This enables these classic optimization algorithms to break free from the blindness of exploration and achieve faster convergence and better results, demonstrating the wide applicability and great potential of this framework in improving the performance of tools in the entire field of computational protein design.

## Introduction

Protein design aims to discover high-fitness sequences with tailored functional attributes, such as enzymatic activity, thermodynamic stability, and binding affinity [1, 2], by exploring astronomically vast fitness landscapes. The directed evolution (DE) framework is a cornerstone of protein sequence optimization through iterative cycles of random mutagenesis, high-throughput screening, and selective pressure to optimize sequences [3, 4]. While DE has enabled breakthroughs in enzyme engineering [5] and antibody design [6], it is constrained by its dependence on experimental screening. This limitation is further exacerbated by the combinatorial explosion of sequence space, which makes exhaustive search infeasible. For instance, a 100-residue protein corresponds to a design space of 20^100^ (∼10^130^) possible variants. As fitness requirements increase, functional sequences become exponentially rarer [7], necessitating computational strategies to mitigate combinatorial explosion.

To address these challenges, various computational strategies based on evolutionary algorithms have been proposed. Genetic algorithms (GAs) [8] simulate evolutionary processes through mutation, crossover, and selection. Early applications, such as GA-driven synthesis of bioactive compounds [9] and directed evolution of hexapeptides [10], demonstrated efficiency of evolutionary algorithms via iterative optimization. However, GAs struggle with diversity and efficiency in long-sequence optimization [9, 10] due to random mutagenesis in the large combinatorial space, which contains vast invalid sequences. Hybrid machine learning (ML)-evolutionary approaches partially address this by guiding mutations to reduce blind exploration. For instance, Yoshida *et al*. [11] combined a generalized linear model with GA to predict the activity effects of amino acid substitutions, enabling guided mutations in antimicrobial peptide optimization. Zhang *et al*. [12] narrowed the mutation space by integrating multiple algorithms (FCM clustering, PLSR regression, Monte Carlo simulation), and subsequently screened high-frequency mutations to obtain highly active antimicrobial peptides. Nevertheless, the improvement of optimized sequences was highly dependent on the quality of natural peptides, and scalability to long sequences or complex sequences remains limited. In order to further strengthen the exploration of innovative sequences and reduce dependence on natural sequences, EvoPlay framework [13] employed Monte Carlo Tree Search (MCTS), taking single-point mutation action and policy value neural network to guide protein sequence optimization. PMMG [14] introduce Pareto-optimized multi-property design, enabling the generation of diverse protein sequences. These GAs and MCTS-based approaches lack context-aware mutation guidance, leading to blindness and inefficiency of mutation.

Deep learning approaches leverage structural information or functional attributes to seek high-potential mutations [15]. This strategy makes the exploration focus on few important sites, therefore largely reducing the exploration space. For instance, AB-Gen [16] integrated generative pre-trained Transformer (GPT) and deep reinforcement learning (RL) for antibody CDRH3 optimization under multi-attribute constraints. DeepDirect [17] employed adversarial learning to discover affinity mutation sites. MMSite [18] leveraged multimodal fusion for revealing active site identification. ProMEP [19] demonstrated zero-shot mutation effect prediction via sequence-structure representation learning. However, there are still key challenges in efficiently exploring multi-site mutation space, maintaining structural integrity. Moreover, these methods require extensive training data, such as high-quality structural annotations, multi-attribute text descriptions, or protein interaction databases, limiting generalizability to novel targets without task-specific retraining. These limitations highlight the need for flexible, data-efficient models that can operate without relying on explicit structure or large labeled datasets.

Protein language models (ProtLMs), build on Transformer architectures [20], have emerged as revolutionary tools by encoding evolutionary constraints from vast datasets (e.g. UniRef90 [21]). State-of-the-art ProtLMs (e.g., ESM-2 [22], ProtT5 [23]) capture epistasis relationships and co-evolutionary dependencies by masking specific positions and inferring likelihoods of all 20 amino acids. Different with traditional evolutionary algorithms (e.g. GAs and MCTS) that rely on random mutagenesis, ProtLMs offer a principled approach to constrain combinatorial search spaces by heuristically inferring substitutions. This capability provides the foundation for more guided and evolutionarily consistent optimization strategies.

In this study, we proposed ProtHMSO, a heuristic multi-site optimization framework that leverages masked ProtLMs for protein sequence design. ProtHMSO employed substitution probability predicted by ProtLMs to heuristically guide the mutation selection, constraining combinatorial search spaces through biophysical and evolutionary priors. Mutant sequences were iteratively refined through fitness evaluation, with top candidates updating the training set for subsequent optimization cycles. We also integrated ProtHMSO as a mutation operator into GAs and MCTS frameworks to enhance their exploration and convergence efficiency. Experimental results demonstrated ProtHMSO has superiority in generating high-fitness and natural-aligned sequences over conventional evolutionary algorithms. Our contributions are: **(1) A novel heuristic framework:** ProtHMSO leverages masked ProtLM predictions to prioritize substitutions that preserve structural integrity and function, narrowing the exploration space for sequence optimization. **(2) Efficient directed-optimization:** Iterative fitness evaluation and sequence refinement enable rapid convergence to high-performance variants. **(3) A modular plugin:** ProtHMSO enhances GA and MCTS efficiency as a modular plugin, demonstrating broad applicability in protein engineering.

## Preliminary knowledge and Related work

### Protein Language Models

Protein language models (ProtLMs) [24] is capable of effectively capturing the contextual information inherent in protein sequences, demonstrating success applications in structure prediction, function prediction, protein design and evolutionary analysis. Typically, the BERT-style architectures are structured with the encoder component of Transformer [20], which masks parts of the input sequence and predicts the missing residues via self-attention mechanism. This self-attention mechanism can be mathematically represented as:

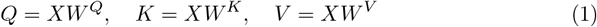

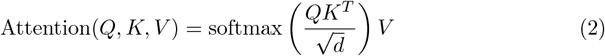

where *X* ∈ ℝ^*L*×*d*^ is the embedded representation of input sequence, which will be transformed into the Query, Key, and Value vectors *Q, K, V*. The matrices *W*^*Q*^, *W*^*K*^, *W*^*V*^ are learnable parameters. Language models commonly use multi-head attention mechanisms to obtain richer information, represented as:

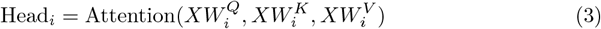

where 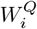, 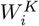, 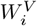 are learnable parameters. The output of multi-head attention is the concatenation of all attention heads, followed by a linear transformation:

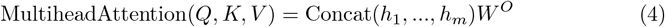

where *h*_*i*_ is the representation of Head_*i*_, and *W*^*O*^ is a learnable parameter.

The pre-trained protein language model, such as ESM-2 [22], utilizes a modified BERT [24] architecture, which consists solely of an encoder and incorporates rotary position embeddings (RoPE) [25] to facilitate inference beyond its trained context window. ESM-2 harnesses the power of the language model to predict the atomic-level structure of proteins, achieving enhanced accuracy and resolution by significantly expanding its parameters, from 8 million to 15 billion. Unlike traditional methods that depend on multiple sequence alignment (MSA), ESM-2 directly infers the 3D structure of proteins from amino acid sequences. This approach dramatically accelerates the rate of structural prediction, making it up to 60 times faster than current state-of-the-art technologies such as AlphaFold [26] and RosettaFold [27]. ESM-2 has been employed in MLDE framework [28] for mutation optimization of sequence iterations. Comparative evaluations across eight benchmark datasets demonstrated that ESM-2 outperforms state-of-the-art methods, highlighting the validity of utilizing the ESM-2 model for sequence mutation. In our ProtHMSO framework, ESM-2 was also employed to predict sequence mutations, followed by the direct application of a pre-trained fitness scorer, eliminating the need for iterative optimization to achieve optimal results.

### Genetic Algorithm

The Genetic Algorithm (GA) serves as a search method designed to solve optimization problems. Its fundamental principle was inspired by the mechanisms of natural selection and genetics in biological systems. The following outlines the key principles and components of genetic algorithms:

- **Natural selection**: Natural selection is a core mechanism of biological evolution. In genetic algorithms, this principle is used to determine which individuals will be selected for reproduction based on the evaluation of their fitness. Individuals with higher fitness have a greater likelihood of being selected, thereby progressively optimizing the quality of the solution.
- **Genetics**: The genetic principles are incorporated into genetic algorithms through the operations of crossover and mutation. The crossover operation simulates the recombination of genetic material, while the mutation operation introduces random alterations in genes. Together, these operations generate new individuals, enhance population diversity, and explore new regions in the solution space.

GAs are fundamentally an efficient, parallel, and global search method that can autonomously acquire and accumulate knowledge about the search space throughout the process. They also adaptively control the search to identify the optimal solution. The application of genetic algorithms to optimize protein sequences is a widely used optimization approach. In this process, a protein sequence dataset is employed to initialize the population, which is then subjected to hybridization or mutation to produce offspring. The offspring are subsequently evaluated using a fitness function to form a new population, and the process is iteratively evolved. For instance, the integration of machine learning with genetic algorithms has been applied to optimize protein sequences [11], [29], where genetic algorithms, along with experimental feature feedback, enable the optimization of molecules with desired biological or chemical activity starting from a small initial library [30]. While these methods have demonstrated promising predictive capabilities, they typically require relatively large datasets for accurate predictions and precise parameter tuning based on the initial protein sequence library. In our approach, we embedded the ProtHMSO framework as a component within GAs, significantly enhancing the performance of the sequences generated by the algorithm.

### Monte Carlo Tree Search

Monte Carlo Tree Search (MCTS) [31] describes a heuristic search strategy that evaluates the advantages and disadvantages of potential decisions through simulation (or “sampling”) to select the best subsequent action. The basic steps of MCTS include selection, expansion, simulation, and backpropagation. Starting from the root node, the algorithm proceeds according to a specific strategy, such as the Upper Confidence Bounds (UCB) [32, 33] formula, which is a commonly used selection strategy, defined as follows:

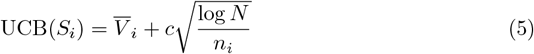

where 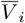 represents the average estimated value of action *i*; *c* is the exploration factor, typically set to 2; *N* is the total number of actions; *n*_*i*_ is the number of action *i* has been taken.

The core idea of MCTS is to balance exploration and exploitation, and the UCB formula plays a crucial role in achieving this balance. It combines the average return, the number of visits to a node, and parameters that control the depth of exploration. This allows MCTS to effectively explore potentially high-quality paths in the search tree. Using the UCB formula, the UCB values of all child nodes can be calculated, and the most promising child node is selected based on the UCB value. This process continues until an unexplored node or a terminal node is reached.

In the expansion step, if the currently selected node is not a terminal node, one or more child nodes are created, representing possible next actions. These newly created child nodes are unexplored, so a random child node is selected to simulate the process until the game ends or a predetermined simulation depth is reached. The results of the simulation are then back-propagated along the selected path, updating the statistics of the nodes along the way, such as the number of visits. MCTS has been successfully applied to the optimization of protein sequences, as demonstrated by the EvoPlay framework proposed by Wang et al. [13]. This self-play reinforcement learning framework, based on the single-player version of AlphaZero [34], interacts with a policy-value neural network and a look-ahead MCTS to optimize sequences with single-point mutations, guided by both breadth and depth. In our work, we integrated ProtHMSO as a component within MCTS to enhance its effectiveness.

## Methodology

In this section, we present ProtHMSO, a heuristic multi-site optimization strategy. First, we introduced the core algorithm of ProtHMSO, which integrates a masked ProtLM to predict beneficial mutations as the heuristic search strategy. Then, we show how ProtHMSO can function not only as a standalone framework but also as a plug-and-play module that improves the convergence of genetic algorithms (GA) and Monte Carlo tree search (MCTS).

The goal of directed evolution is to find the optimal sequence *X*^*\**^ that maximizes a particular objective function:

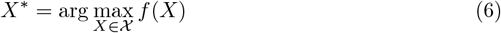

where χ represents the mutation space of sequences, and *f* (*X*) is the fitness function of sequence *X* in χ. During optimization, the algorithm selects the best mutant sequence from those generated by mutations such that the returned fitness *y*^*\**^ = *f* (*X*^*\**^) is the optimal fitness among all *y* values.

### Heuristic Multi-site Optimization Framework

Heuristic multi-site optimization (HMSO) framework leverages the predicted beneficial mutations by ProtLMs for context-aware sequence exploration. The core mechanism of ProtLMs is masked language modeling (MLM), which is unsupervised pretraining strategy by predicting masked tokens in varied contexts, therefore enabling dynamic context-aware mutation effects. Given a target sequence, a ProtLM is employed to predict the likelihood of substituting a specific residue with one of the 20 amino acids, thereby quantifying the impact of mutations on sequence fitness [35]. Specifically, a protein *X* = (*x*_1_, …, *x*_*i*_, …, *x*_*L*_) has *L* residues, where the set of masked positions is *M* = {*m*_1_, …, *m*_|*M*|_}. The masked residues can be inferred based on the surrounding contextual information. The probability of masked tokens is computed as Π_*m∈M*_ *p*(*x*_*m*_ | *X*_*\M*_), where *X*_*\M*_ means all input tokens except those masked tokens. In this study, ESM-2 [22] was primarily used as the masked ProtLMs to predict mutation sites of input sequences. The process of HMSO framework is illustrated in **Fig 1**.

#### Algorithm 1 Heuristic Multi-site Optimization Framework

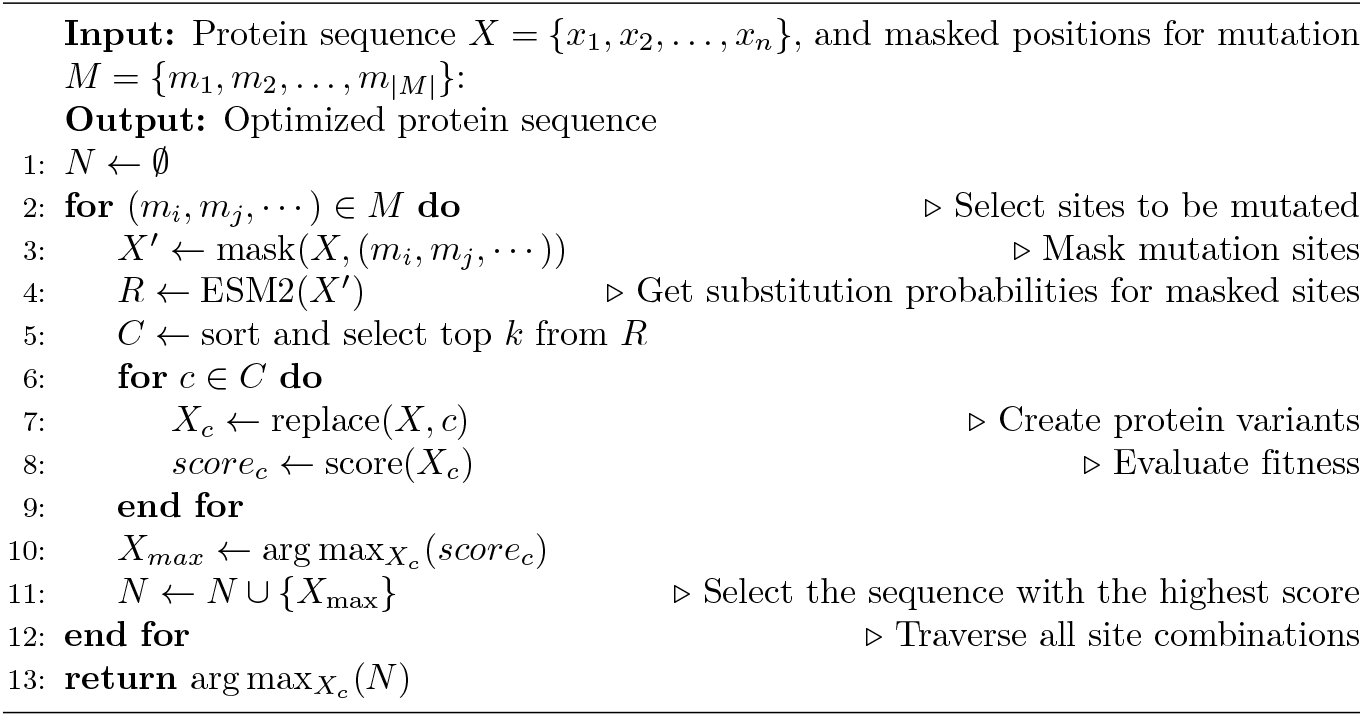

**Fig 1.**
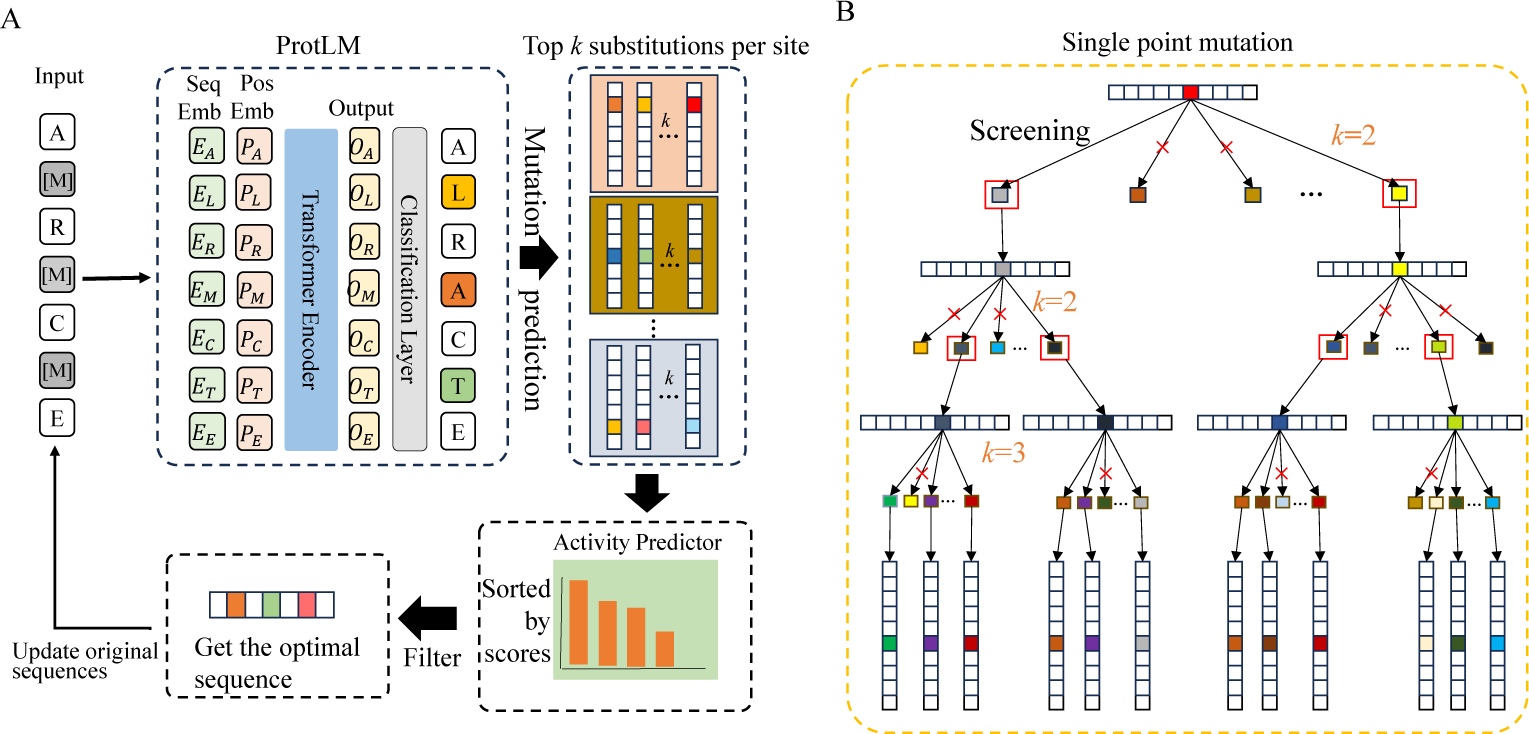
Illustration of heuristic multi-site optimization for protein sequence design using masked protein language models. (A) The iterative process of multi-site mutagenesis using ProtHMSO. (B) A illustration of narrowing exploration space via single-site mutation.

The core process of our proposed HMSO is described in **Algorithm 1**. HMSO takes a protein sequence *X* and the corresponding mutation sites *M* as inputs, and outputs the optimized sequence *N*. The protein sequence *X* is first masked according to the selected mutation sites and then fed into the ESM-2 model to get their substitution probabilities of 20 natural amino acids at each mutation site. These probabilities are then sorted in descending order, the top *k* substitutions are selected, and the corresponding *k* protein variants are generated. Then these *k* variants are assigned a score by a fitness evaluator, subsequently the sequence with the highest score in each mutation is selected and added to the final set of mutant sequences. This process is iterated to eventually obtain an optimized sequence.

### Heuristic Genetic Algorithm based on HMSO

HMSO is not only a standalone framework but also a modular plugin that enhances the convergence efficiency of genetic algorithm (GA) by guiding exploration with evolutionary priors. In traditional GAs, random crossover/mutation operators lack biological plausibility, often disrupt conserved structural motifs, and single-site optimization neglects epistatic interactions between distant residues. To overcome these challenges, we proposed GA-HMSO, a heuristic genetic algorithm based on HMSO, by employing HMSO to heuristically guide GA mutation operation, enabling it to address the limitations of conventional random mutation and low-diversity offspring generation. First, we initialize the population with the prepared protein sequence, and then select mutation and crossover operations based on probability. The crossover process uses two-point crossover, and the mutation process uses ESM-2 to guide mutation. After completing the crossover mutation operation, the generated sequence is evaluated, and then the offspring is screened according to the fitness function to update the parent generation. This heuristic framework enables simultaneous optimization of multiple sites, leveraging the ProtLMs ability to capture long-range interactions and evolutionary constraints, which is particularly critical for navigating complex fitness landscapes in protein design. At the same time, we also use dynamic crossover mutation probability to prevent it from falling into the local optimal solution problem. The framework of GA-HMSO is illustrated in **Fig 2**.

#### Algorithm 2 Heuristic Genetic Algorithm based on HMSO

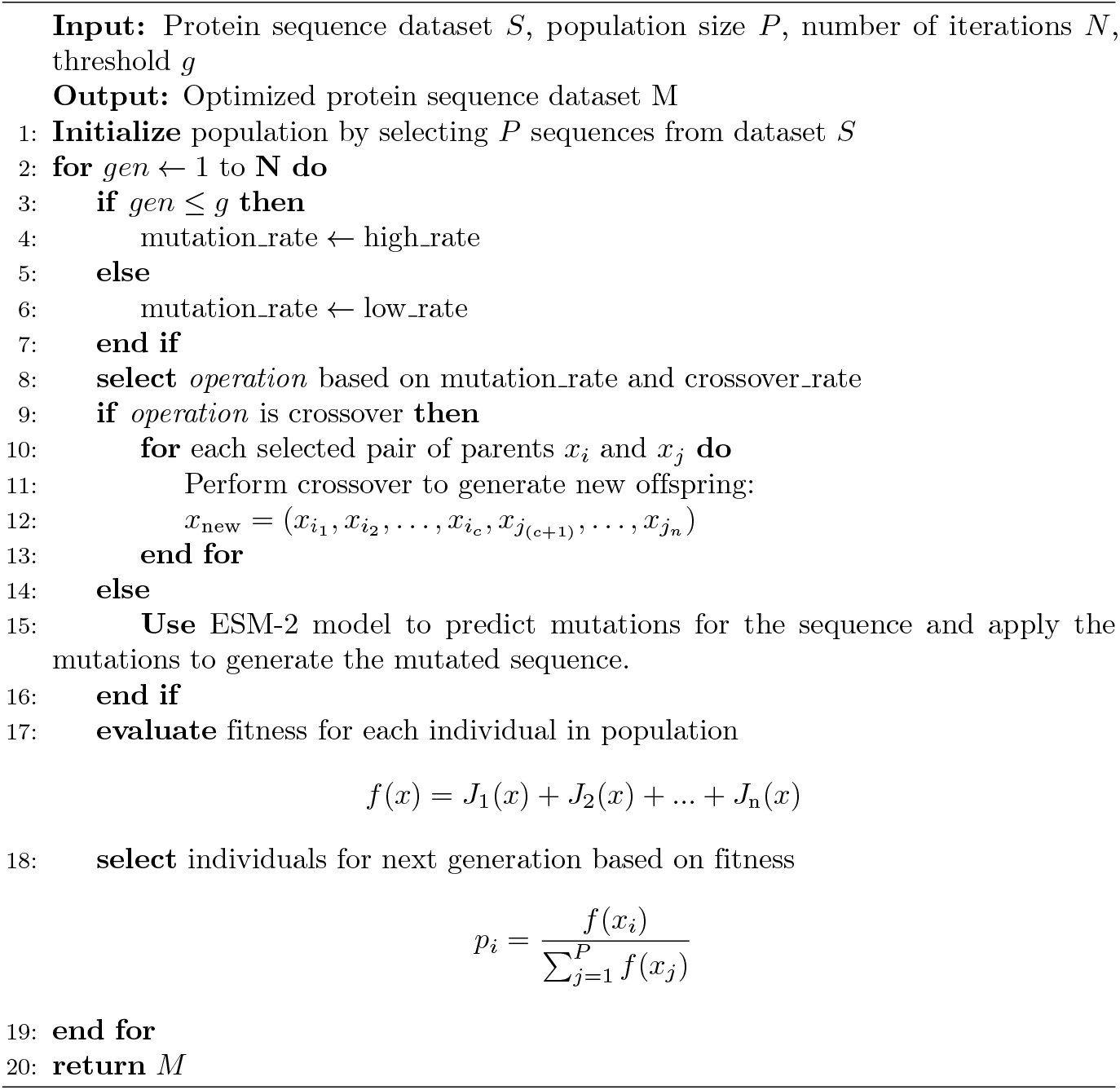

**Fig 2.**
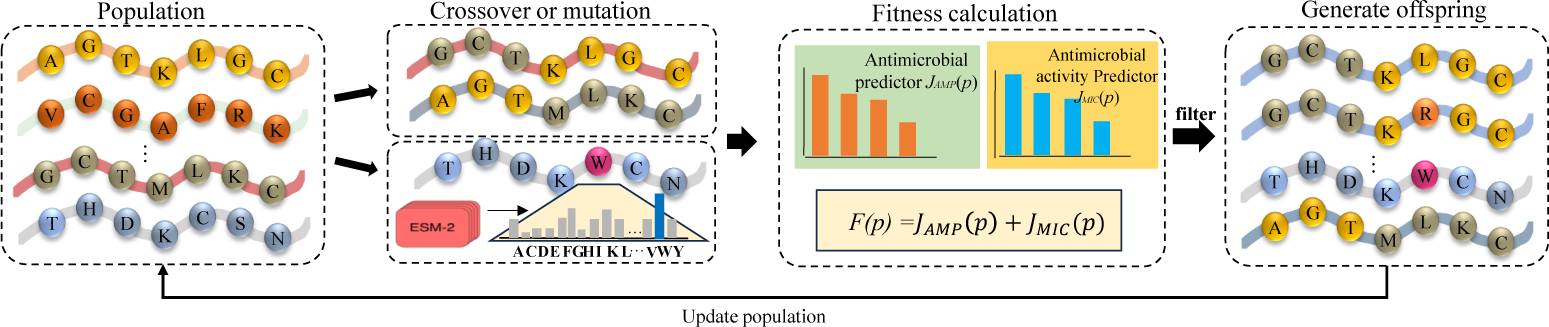
Improved genetic algorithm framework using ESM-2. The protein sequence is used to initialize the population. In the mutation operation, the ESM-2 model is used instead of random mutation. The weighted scores of each predictor and the custom error term are used as the fitness score to screen the offspring and update the population.

An individual being selected into the next population is depending on its fitness. Let *p*_*i*_ be the probability of the *i*-th individual being selected, then:

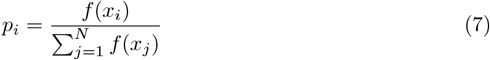

where *f* (*x*_*i*_) represents the fitness of the *i*-th individual, and *N* denotes the number of individuals in the generated offspring. The definition of the fitness function *f* (*x*) is a combination of multiple attributes, as follows:

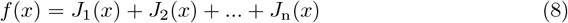

where *J*_i_(*x*) represents the attribute score of the protein sequence. The scores of these predictors are added together as the final score. Therefore, GA-HMSO can be employed for multiple objective optimization. Selected individuals are then combined through a crossover operation to generate new offspring. Assuming the crossover point is *c*, and considering two individuals *x*_*i*_ and *x*_*j*_, the offspring *x*_new_ can be represented as *x*_new_ = (*x*_*i*1_, *x*_*i*2_, …, *x*_*ic*_, *x*_*j*(_, …, *x*_*jn*_).

During the selection phase, we use the ‘[mask]’ token to replace the amino acids at selected positions of the sequences, represented by *x*_*m*_ = (*x*_1_, *x*_2_, *· · ·*, [*mask*], *· · ·*, [*mask*], *· · ·, x*_*n*_). This mask vector allows for the selective and dynamic combination of sequence components, thereby enhancing the algorithm’s efficiency in optimization. In the early iterations, we set a high mutation probability to promote exploration. As the number of iterations increases, the mutation probability decreases to refine the local optimization. Specifically, the mutation probability *P*_MUT_ and the crossover probability *P*_CX_ which were manually chosen vary according to the generation number *gen* as follows:

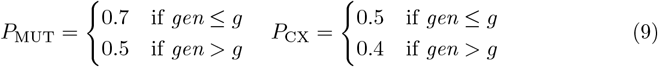

where *P*_MUT_ represents the mutation probability, and *P*_CX_ represents the crossover probability, with *g* being the generation threshold after which the probabilities are adjusted. This adjustment is due to the fact that a higher mutation probability enhances global optimization, but after a certain number of generations, the population tends to converge to local extrema. At this point, with a large number of mutations, the mutation probability is reduced in the later stages to focus more on local optimization. Finally, offspring are selected based on their fitness scores, and the selected offspring are used to update the population for the next iteration. The process of GA-HMSO is described in **Algorithm 2**.

### Heuristic Monte Carlo Tree Search based on HMSO

Traditional Monte Carlo Tree Search (MCTS) relies on random simulation and node selection strategies based on UCB formula. It is difficult to efficiently explore potential mutation combinations in the high-dimensional space of protein sequence design. In its expansion step, the number of child nodes generated by each node grows exponentially with the length of the sequence, resulting in a shallow search tree, a large number of valuable mutation paths being ignored, and falling into a local optimal solution. For example, in the design of antimicrobial peptides, traditional MCTS requires thousands of iterations to converge, and often generates invalid sequences due to random mutations, wasting computing resources. To address this, in this study, we proposed MCTS-HMSO by combining MCTS with HMSO to heuristically optimize protein sequences. The framework is shown in **Fig 3**.

**Fig 3.**
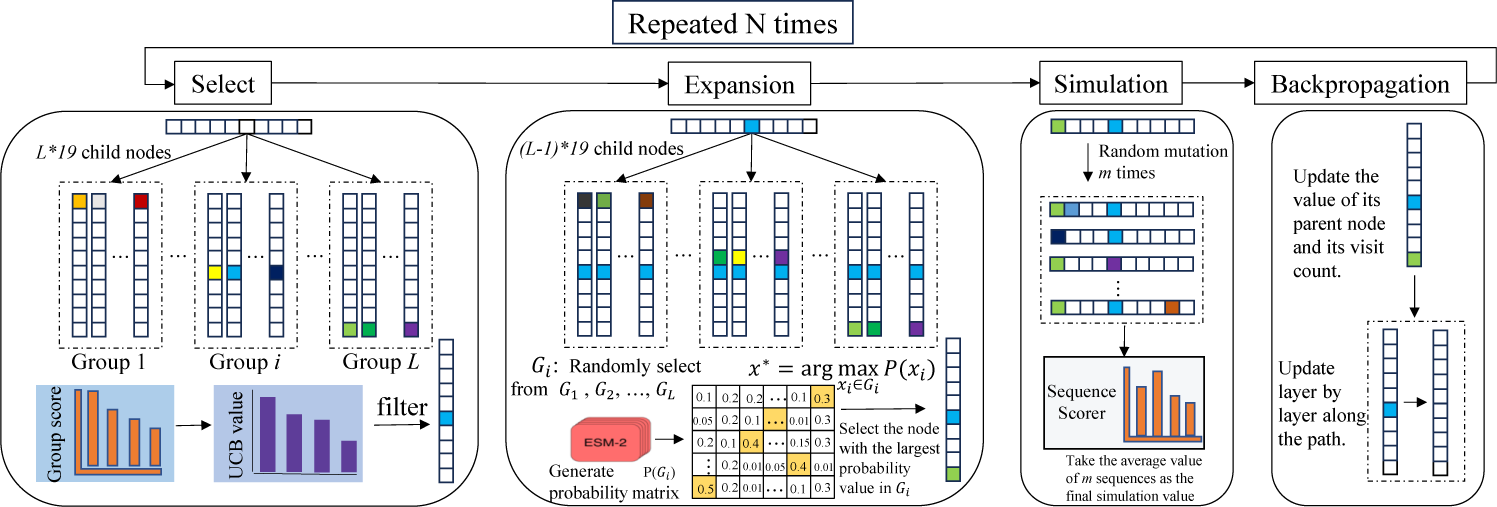
Heuristic Monte Carlo Tree Search based on HMSO. First, the subnode to be expanded is selected according to the UCB value, and then the top *k* subnodes are generated for each site according to the probability matrix generated by ESM-2. Finally, one of the subnodes is randomly returned. After simulating the subnodes, they are updated layer by layer along the generation path of the subnodes.

In MCTS-HMSO, a node represents a protein sequence. The root node is the original protein sequence to be optimized. The structure of the entire tree is as follows: the first layer contains the root node, and the second layer contains 19 *× L* child nodes, each representing one of the 19 *× L* possible mutated sequences. When the child nodes of the second layer are expanded, each generates 19 *×* (*L* — 1) child nodes, meaning that previously mutated positions will no longer undergo mutation in subsequent layers. Instead, mutations will occur at different positions. Generally, the number of child nodes generated at the *i*-th layer is 19 *×* (*L* –*i* –1) (with the root node being the first layer).

During the expansion step, to heuristically explore these potential protein variants, all sub-nodes are grouped into *L* groups according to their mutation positions. Then, each sub-node is assigned a substitution score by employing ESM-2. Each group is also assigned a score by summing the UCB values of all nodes within the group. The initial score of each group is set to infinity, and the initial UCB value of each sub-node is also infinite.

In the selection process, the group is first selected based on the score of each group, and then the sub-node with the highest UCB value is chosen within the selected group. Specifically, given a group *G*_*i*_ consisting of nodes *x*_1_, *x*_2_, …, *x*_*n*_, where each node *x*_*j*_ has a UCB value *UCB*(*x*_*j*_). The nodes that have been explored are *x*_1_, *x*_2_, …, *x*_*e*_ and the score of the group *S*(*G*_*i*_) can be represented as:

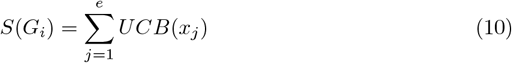

where *e* is the number of nodes explored in group *G*_*i*_, and the *UCB* value of each node is still calculated according to Eq. (5). After calculating the *UCB* value, the score of the group to which the current node belongs will be updated accordingly. Finally, the final child node is selected according to the following formula:

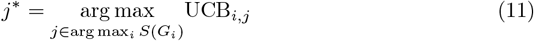

This sequence is then used for the next simulation step. In our experiment, we perform a series of random mutations on the selected sequence, and then evaluate the fitness of mutated variants, at last take the average of fitness scores as the final simulated value for the selected sequence. During the backtracking process, both the sub-node score and its group score need to be updated. The algorithm of MCTS-HMSO is outlined in **Algorithm 3**.

#### Algorithm 3 Heuristic Monte Carlo Tree Search based on HMSO

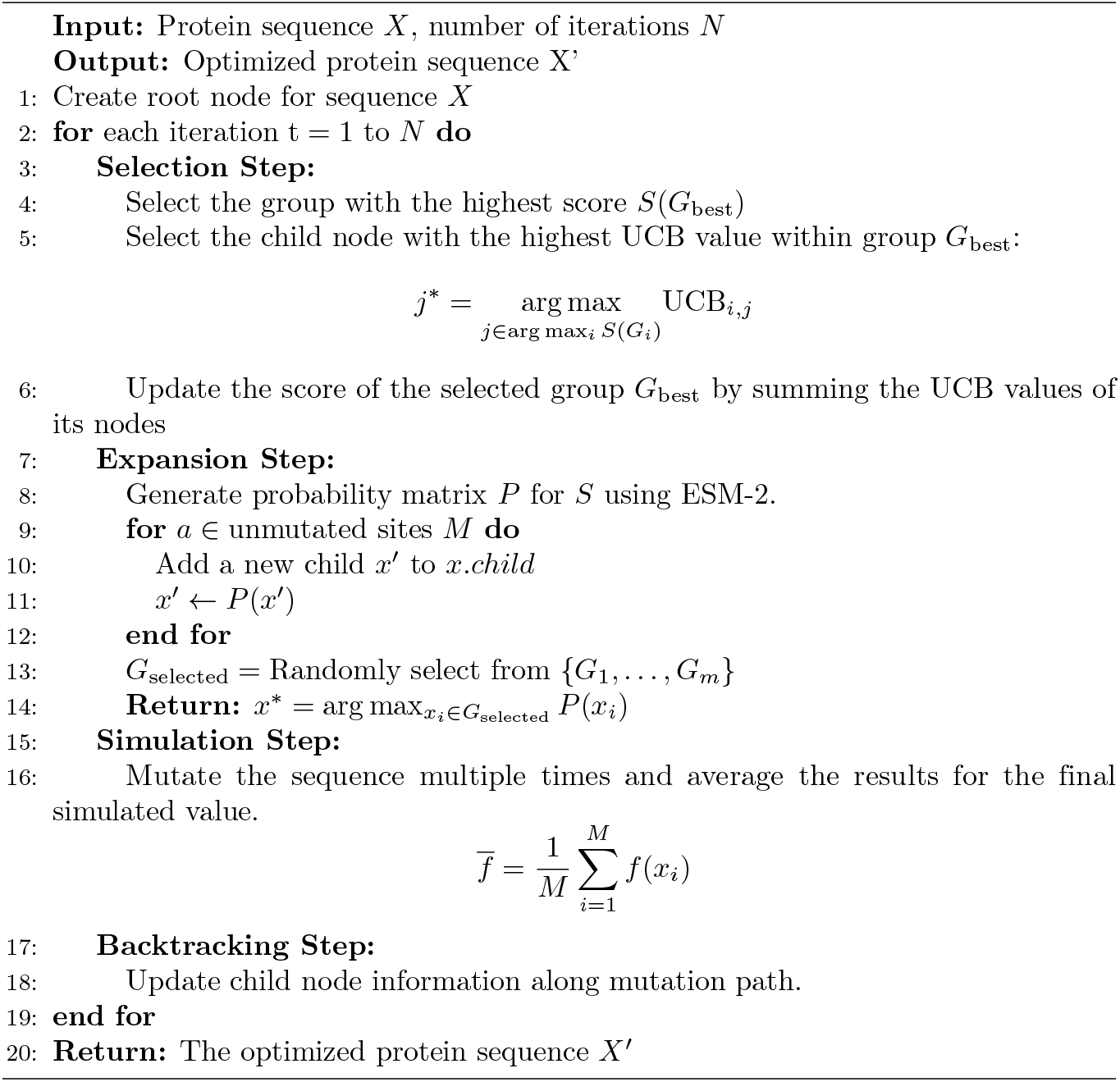

## Experiments

To evaluate the efficacy of ProtHMSO, we conducted comparative analyses against random mutation strategies, conventional GA and conventional MCTS. These comparisons aimed to assess both the stand-alone performance of ProtHMSO and its integration as a heuristic component in GA (GA-HMSO) and MCTS (MCTS-HMSO). Experiments were performed on two benchmark datasets: an AMP dataset and the ProteinGym database, enabling comprehensive evaluation across diverse optimization scenarios.

### Masked protein language models

We employed ESM-2 (650M), as the masked ProtLMs, to heuristically guide mutation. The ESM-2 architecture has 650M parameters, including 33 transformer layers with a maximum sequence length of 1,024 tokens, a hidden dimension of 1,280 and 20 attention heads. Its vocabulary includes 25 amino acids, including 5 non-natural residues, while the predictor can only recognize the 20 natural amino acids. To ensure biological relevance, predictions corresponding to non-natural amino acids were substituted with lysine (K), following the protocol established by Zhang et al. [36]. All optimizations were performed using single-precision floating-point arithmetic on NVIDIA A100 GPUs.

### Antimicrobial Peptide Dataset

All antimicrobial peptides (AMPs) were sourced from the Database of Antimicrobial Activity and Structure of Peptides (DBAASP) [37], which catalogs peptides along with activity values and known hemolytic behaviors. From the initial dataset comprising 11,805 non-cyclic peptides, we retained 4,505 unique active sequences after deduplication, following the preprocessing by Capecchi *et al*. [38]. To systematically evaluate the performance of ProtHMSO across diverse optimization scenarios, sequences were stratified into four disjoint subsets based on the antimicrobial likelihood *P*_AMP_ and activity potential *P*_MIC_: **Case 0: High-active AMPs** (*P*_AMP_ ≥ 0.8, *P*_MIC_ ≥ 0.8).

These sequences already exhibit optimal antimicrobial properties and do not require further optimization. **Case 1: Activity-enhancement candidates** (*P*_AMP_ ≥ 0.8, *P*_MIC_ ≤ 0.8). Optimization targets improvement of antimicrobial activity while preserving functionality. **Case 2: Functionality-enhancement candidates** (*P*_AMP_ ≤ 0.8, *P*_MIC_ ≥ 0.8). Optimization focuses on enhancing the antimicrobial function. **Case 3: High-challenge sequences** (*P*_AMP_ ≤ 0.8, ≤ *P*_MIC_ 0.8). These sequences represent the most challenging optimization scenario and demand optimization of both functionality and activity. There are 875, 184, 476 peptide sequences in Case 1, Case2 and Case3.

### ProteinGym Datasets

ProteinGym [39] is a collection of benchmarks aiming at comparing the ability of models to predict the effects of protein mutations. The benchmarks in ProteinGym were divided according to mutation type (substitutions vs. indels) and ground truth source (DMS assay vs. clinical annotation). In this study, we focused on the substitution type, where DMS benchmark contains 217 assays spanning over 2.4 million mutants and clinical benchmark contains 2,525 genes spanning over 63 thousand mutants. The mutant fitness of DMS substitutions were evaluated by DMS score, which is a continuous experimental measurement. Higher values indicate high-fitness, while the mutant fitness of clinical substitutions were evaluated by DMS score bin, which is a binary label indicating whether the DMS score exceeds a fitness threshold (1: pathogenic, 0: benign).

To further investigate site-specific optimization, we conducted targeted mutagenesis at functional residues. Active sites were first identified using NCBI’s Conserved Domain Database (CDD) and literature annotations. Mutants were then introduced at these positions via both random and ProtHMSO methods. The resulting mutants were assessed by comparing their DMS score distributions to quantify performance differences between the two strategies.

### Metrics

To holistically evaluate our heuristic optimization framework, we employed four complementary metrics spanning functional and structural assessments.

For the AMP optimization, two functional fitness metrics: antimicrobial probability (*P*_AMP_ ∈ [0, 1]) and minimal inhibitory concentration probability (*P*_MIC_ ∈ [0, 1]) were employed to quantify the likelihood of a sequence being antimicrobial peptide and retaining effective antimicrobial activity against gut microbiota at low concentrations (≤ 32 *µ*g/mL). These indicators were predicted by the classification model derived from HydrAMP [40].

For the evaluation of structural validity and biological plausibility of designed sequences, we utilized two additional indicators: predicted local Distance Difference Test (pLDDT) scores and perplexity. pLDDT scores reflect the confidence in predicted backbone atom positions with higher values indicating greater structural validity. In this study, pLDDT scores were computed via ESMFold [41]. Perplexity evaluates the negative log-likelihood of sequences under a generative protein language model. Lower perplexity values indicate closer alignment with natural sequence distributions, correlating with higher experimental expressibility. In this study, perplexity is calculated via ProGen2 [42].

## Results and discussion

We comprehensively evaluate ProtHMSO across multiple experimental configurations, benchmarking its efficacy with random mutation strategies, and then assessing its integration with GAs and MCTS.

### Improving functionality and activity of antimicrobial peptides via ProtHMSO

We first compared ProtHMSO with random mutagenesis on the AMP dataset. Random mutation is a mutation with equal probability at each site. Since peptides are short amino acid sequences with relative same mutation space, we could conduct multiple mutations simultaneously. Mutants were performed across 1-5 variable sites per sequence. Due to the vast compositions, we limited the mutation times to reduce exploration space. For single-point mutations, each site of the sequence was mutated once across the sequence length, and the variant with highest score was selected as the final optimized sequence. For multiple-point mutagenesis, the mutated sites in each iteration were first randomly selected, and then all substitutions were determined simultaneously via random mutagenesis or ProtHMSO, respectively. The number of mutants is the length of the corresponding sequence. The variant with the highest score is selected as the final optimized sequence of the corresponding mutation method.

The evaluation of optimized sequences on the three cases in AMP dataset in terms of the average *P*_AMP_ and *P*_*MIC*_ (**Table 1**) and their distributions (**Fig 4**) showed that both random and ProtHMSO methods enhanced the activity and functionality of all three cases, compared with the original sequences across 1-5 sites mutation. Moreover, ProtHMSO consistently performed better than random method, where ProtHMSO achieved the highest average *P*_MIC_ of 0.82 for Case1, *P*_AMP_ of 0.96 for Case2, and *P*_AMP_ of 0.86 and *P*_MIC_ of 0.56 for Case3. These results demonstrated ProtHMSO can improve functionality and activity of AMPs by heuristically searching, even for the high-challenge candidates that demand optimization of both functionality and activity.

**Table 1.** Comparison the average performance of sequences generated using different mutation methods in terms of *P*_AMP_, *P*_MIC_, pLDDT and perplexity across three AMP cases.

| Methods | Case1 |  |  | Case2 |  |  | Case3 |  |  |  |
| --- | --- | --- | --- | --- | --- | --- | --- | --- | --- | --- |
| | $P_{MIC} \uparrow$ | pLDDT↑ | perplexity↓ | $P_{AMP} \uparrow$ | pLDDT↑ | perplexity↓ | $P_{AMP} \uparrow$ | $P_{MIC} \uparrow$ | pLDDT↑ | perplexity↓ |
| Random_M1 | 0.57($\pm 0.42$) | 70.51($\pm 11.68$) | 17.45($\pm 5.23$) | 0.85($\pm 0.24$) | 70.82($\pm 8.39$) | 17.65($\pm 3.53$) | 0.68($\pm 0.34$) | 0.26($\pm 0.39$) | 67.89($\pm 11.55$) | 17.21($\pm 4.59$) |
| Random_M2 | 0.63($\pm 0.42$) | 70.01($\pm 11.45$) | 18.21($\pm 4.91$) | 0.87($\pm 0.22$) | 70.81($\pm 9.00$) | 18.16($\pm 3.55$) | 0.74($\pm 0.32$) | 0.30($\pm 0.41$) | 67.69($\pm 11.33$) | 17.85($\pm 4.58$) |
| Random_M3 | 0.66($\pm 0.43$) | 69.34($\pm 11.26$) | 18.72($\pm 4.95$) | 0.89($\pm 0.23$) | 70.75($\pm 8.58$) | 18.85($\pm 4.69$) | 0.78($\pm 0.30$) | 0.37($\pm 0.45$) | 67.09($\pm 11.19$) | 18.37($\pm 4.51$) |
| Random_M4 | 0.67($\pm 0.42$) | 68.56($\pm 11.16$) | 19.33($\pm 4.71$) | 0.89($\pm 0.14$) | 70.12($\pm 9.11$) | 19.22($\pm 3.84$) | 0.82($\pm 0.27$) | 0.38($\pm 0.45$) | 66.72($\pm 11.49$) | 18.70($\pm 3.91$) |
| Random_M5 | 0.65($\pm 0.43$) | 67.67($\pm 11.00$) | 19.85($\pm 4.54$) | 0.91($\pm 0.20$) | 70.02($\pm 8.71$) | 19.63($\pm 3.31$) | 0.84($\pm 0.25$) | 0.42($\pm 0.46$) | 65.98($\pm 11.05$) | 19.39($\pm 4.20$) |
| ProtHMSO_M1 | 0.58($\pm 0.42$) | 71.90($\pm 11.82$) | 15.93($\pm 4.66$) | 0.85($\pm 0.25$) | 73.05($\pm 8.21$) | 16.23($\pm 3.07$) | 0.64($\pm 0.36$) | 0.28($\pm 0.41$) | 69.73($\pm 11.46$) | 16.08($\pm 4.29$) |
| ProtHMSO_M2 | 0.67($\pm 0.41$) | 71.84($\pm 11.76$) | 15.86($\pm 4.35$) | 0.89($\pm 0.22$) | 73.56($\pm 8.73$) | 15.99($\pm 3.15$) | 0.72($\pm 0.34$) | 0.34($\pm 0.44$) | 69.76($\pm 11.52$) | 16.07($\pm 3.76$) |
| ProtHMSO_M3 | 0.76($\pm 0.37$) | 71.75($\pm 11.41$) | 15.66($\pm 4.09$) | 0.93($\pm 0.18$) | 73.92($\pm 8.17$) | 15.56($\pm 2.83$) | 0.79($\pm 0.31$) | 0.45($\pm 0.46$) | 70.24($\pm 11.49$) | 15.80($\pm 3.66$) |
| ProtHMSO_M4 | 0.81($\pm 0.33$) | 71.73($\pm 11.58$) | 15.48($\pm 4.69$) | 0.95($\pm 0.16$) | 73.85($\pm 8.91$) | 15.17($\pm 2.79$) | 0.83($\pm 0.28$) | 0.49($\pm 0.46$) | 71.15($\pm 11.22$) | 15.60($\pm 3.37$) |
| ProtHMSO_M5 | **0.82**($\pm 0.33$) | **71.56**($\pm 11.46$) | **14.82**($\pm 3.74$) | **0.96**($\pm 0.13$) | **74.76**($\pm 8.08$) | **14.43**($\pm 2.68$) | **0.86**($\pm 0.26$) | **0.56**($\pm 0.46$) | **71.25**($\pm 11.26$) | **15.13**($\pm 3.24$) |
| GA | 0.09($\pm 0.28$) | 62.62($\pm 9.92$) | 18.96($\pm 3.04$) | 0.31($\pm 0.41$) | 66.50($\pm 8.11$) | 19.53($\pm 2.78$) | 0.43($\pm 0.37$) | 0.06($\pm 0.20$) | 59.88($\pm 9.18$) | 19.19($\pm 2.10$) |
| GA-HMSO_fixed | 0.43($\pm 0.46$) | 69.41($\pm 10.94$) | 12.17($\pm 3.48$) | 0.88($\pm 0.25$) | 74.68($\pm 8.34$) | 9.82($\pm 4.32$) | 0.60($\pm 0.42$) | 0.29($\pm 0.42$) | 67.98($\pm 11.00$) | 12.93($\pm 3.44$) |
| GA-HMSO_dynamic | **0.75**($\pm 0.41$) | **81.62**($\pm 9.18$) | **5.20**($\pm 2.85$) | **0.90**($\pm 0.29$) | **80.87**($\pm 9.12$) | **5.82**($\pm 2.85$) | **0.90**($\pm 0.25$) | **0.83**($\pm 0.34$) | **82.37**($\pm 9.56$) | **4.52**($\pm 2.56$) |
| MCTS_100 | 0.73($\pm 0.40$) | 70.80($\pm 11.50$) | 17.68($\pm 5.43$) | 0.94($\pm 0.14$) | 71.74($\pm 8.46$) | 17.97($\pm 3.61$) | 0.76($\pm 0.31$) | 0.34($\pm 0.44$) | 67.92($\pm 11.59$) | 17.49($\pm 4.40$) |
| MCTS_300 | 0.81($\pm 0.35$) | 70.66($\pm 11.55$) | 18.06($\pm 5.18$) | 0.96($\pm 0.11$) | 71.05($\pm 8.68$) | 17.95($\pm 4.25$) | 0.82($\pm 0.28$) | 0.42($\pm 0.46$) | 68.21($\pm 11.47$) | 17.62($\pm 4.37$) |
| MCTS_500 | 0.86($\pm 0.31$) | 70.42($\pm 11.73$) | 18.40($\pm 5.80$) | 0.98($\pm 0.08$) | 71.48($\pm 8.75$) | 18.42($\pm 5.08$) | 0.87($\pm 0.24$) | 0.42($\pm 0.47$) | 67.92($\pm 11.56$) | 17.86($\pm 5.04$) |
| MCTS-HMSO_100 | 0.74($\pm 0.40$) | 71.93($\pm 11.45$) | 16.39($\pm 4.63$) | 0.96($\pm 0.13$) | 73.21($\pm 8.39$) | 16.41($\pm 3.06$) | 0.82($\pm 0.30$) | 0.40($\pm 0.46$) | 69.48($\pm 11.36$) | 16.26($\pm 4.01$) |
| MCTS-HMSO_300 | 0.83($\pm 0.35$) | 71.90($\pm 11.43$) | 16.45($\pm 4.45$) | 0.97($\pm 0.12$) | 73.49($\pm 8.17$) | 16.22($\pm 3.07$) | 0.92($\pm 0.21$) | 0.58($\pm 0.47$) | 69.93($\pm 11.50$) | 16.32($\pm 3.92$) |
| MCTS-HMSO_500 | **0.88**($\pm 0.29$) | **72.13**($\pm 11.46$) | **16.35**($\pm 4.39$) | **0.98**($\pm 0.09$) | **73.51**($\pm 9.00$) | **16.32**($\pm 3.12$) | **0.96**($\pm 0.13$) | **0.66**($\pm 0.45$) | **69.93**($\pm 11.54$) | **16.31**($\pm 3.98$) |

The bold numbers represent the best performance within each kind of methods. The arrows ( ↑ ) indicate that higher values are better, while ( ↓ ) indicates lower values are better. Values in parentheses ( $\pm \dots$ ) represent the standard deviation.

**Fig 4.**
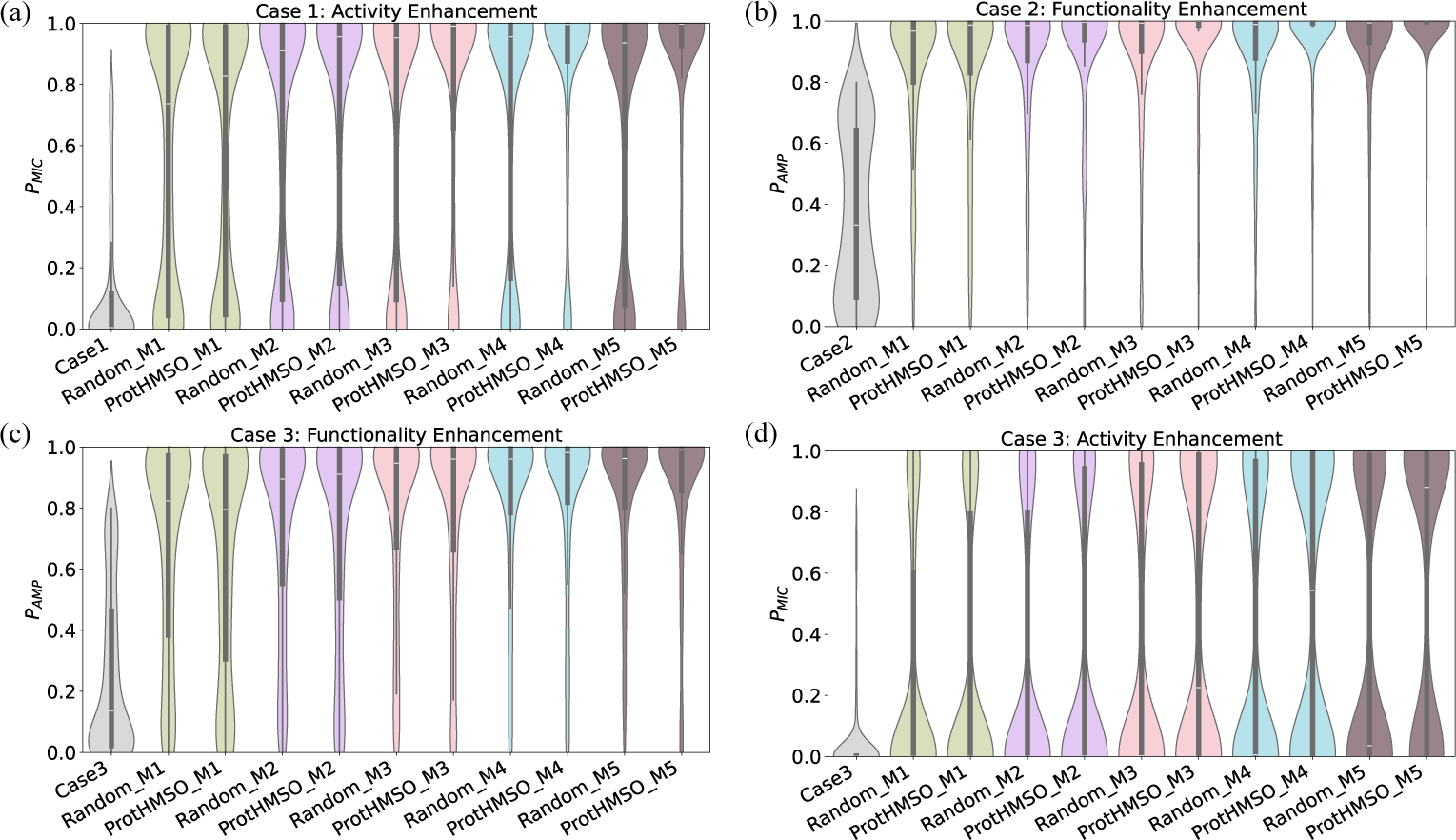
Comparison of functionality or activity enhancement of three AMP cases via random and ProtHMSO guided mutagenesis. (a) The activity-enhancement of AMPs in Case1. (b) The functionality-enhancement of AMPs in Case2. (c-d) The functionality-enhancement (c) and activity-enhancement (d) of AMPs in Case3. ‘Case1’, ‘Case2’ and ‘Case3’ represent the distribution of AMP cases. ‘Random M1’ represents single-site mutation using a random method, and so on.

Besides, theoretically, as the number of mutated sites increases, more potential sequences with high functionality and activity could be explored, subsequently a larger improvement would be achieved. Interestingly, for Case1, random 5-site mutation led to lower activity improvement than 4-site, likely due to exponential sparsity of functional sequences in the combinatorial explosion of mutation space. In the same number of searching iterations, it is exponentially hard to find better variants. However, we observed that ProtHMSO achieved a higher average *P*_MIC_ in the combinatorial explosion of mutation space across all three cases. Notably, as the number of mutated sites increasing, the pLDDT and perplexity metrics achieved by ProtHMSO were also getting better, in contrast the performance of random method was getting worse, indicating that the sequences optimized by ProtHMSO retained high structural validity and sequential plausibility, while the random mutants would break these rationality. This divergence suggests ProtHMSO can leverage epistatic interactions to heuristically guide multi-site combinatorial improvements.

### Exploring beneficial mutation of long protein sequences via ProtHMSO

To rigorously evaluate the efficacy of ProtHMSO, we conducted long protein optimization on the ProteinGym database, where the average length of the Clinical benchmark is 709 and the average length of DMS benchmark is 397. Due to the vast composition, we only focused on the single-site substitutions.

The Clinical benchmark comprises experimentally validated single-site substitutions associated with pathogenicity. We introduced single-site mutations at identical positions in the original sequences using both random mutagenesis and ProtHMSO guided approach. Each sequence generated 10, 50, and 100 mutated variants under each strategy. As shown in **Fig 5**, ProtHMSO significantly outperformed random mutagenesis, generating approximately twice as many non-pathogenic variants as random mutagenesis across all numbers of mutants. These results indicate that ProtHMSO leverages the evolutionary constraints encoded in the protein language model to prioritize functionally benign mutations while mitigating pathogenic outcomes.

**Fig 5.**
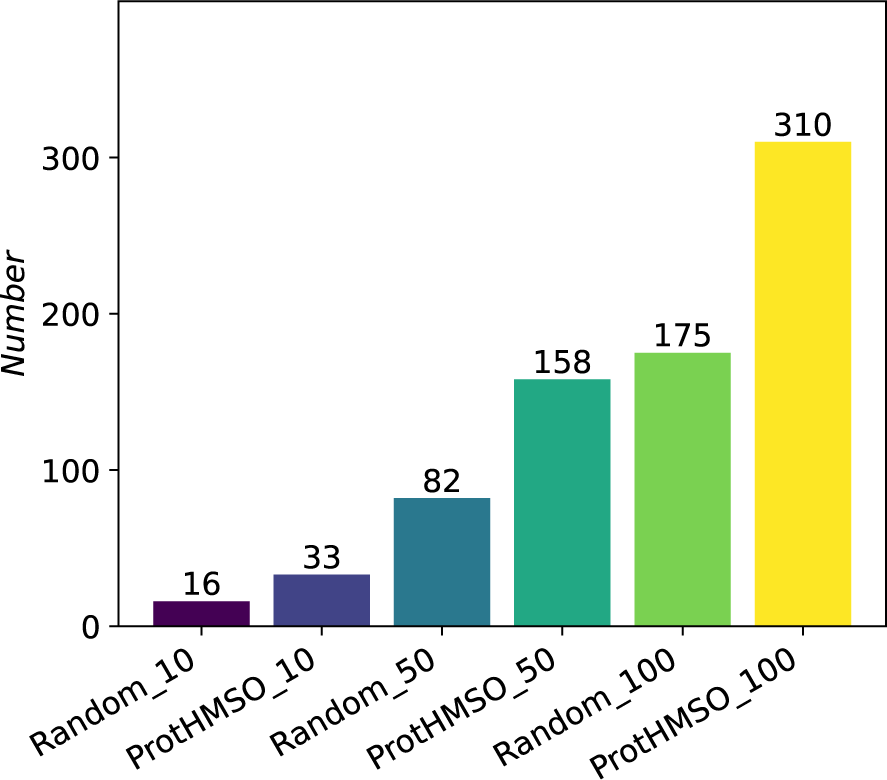
Number of non-pathogenic protein sequences optimized by random and ProtHMSO guided mutagenesis. ‘Random 10’ means to generate 10 sequences using random mutation for each original sequence, and so on.

The DMS benchmark contains both single and two mutant variants with experimentally measured fitness scores (DMS score). We partitioned this dataset into single-site and two-site mutation subsets, then ranking variants within each subset by their DMS score. Functional sites for mutation were identified through conserved domain annotations from the NCBI database, which integrates structural and evolutionary analyses from the Conserved Domain Database [43, 44]. For each sequence, we generated mutant libraries targeting these functional sites using both random mutagenesis and ProtHMSO. The top 10, 20 and 50 experimentally ranked mutants were selected as high-fitness benchmarks. We then assessed the overlap between generated mutants and these top experimental variants. As shown in **Fig 6**, PortHMSO consistently outperformed random mutagenesis across both single and two-site mutations. Notably, ProtHMSO-generated mutants exhibited significantly higher enrichment in top-10, 20 and 50 high-fitness variants. This systematic improvement underscores the advantage of evolutionary-aware language model guidance over stochastic exploration for identifying functional sequence variants.

**Fig 6.**
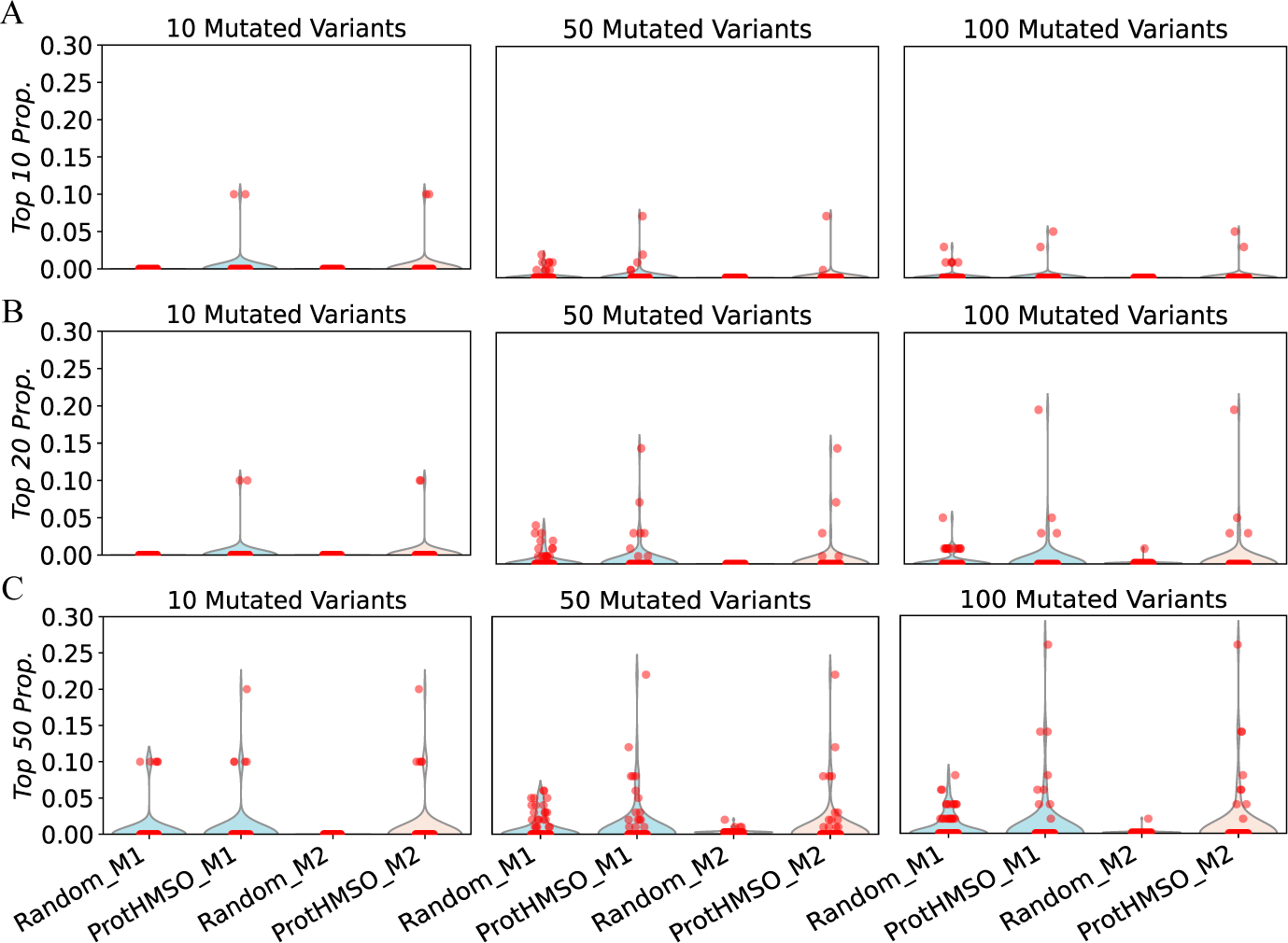
The proportion of mutated variants in top 10, 20 and 50 highest fitness scores. A. The proportion of the top 10 mutated variants with the highest scores among 10, 50, and 100 mutants generated using random mutation and ProtHMSO in the DMS dataset. B. The proportion of the top 20 mutated variants with the highest scores among 10, 50, and 100 mutants generated using random mutation and ProtHMSO in the DMS dataset. C. The proportion of the top 50 mutated variants with the highest scores among 10, 50, and 100 mutants generated using random mutation and ProtHMSO in the DMS dataset. ‘Random_M1’ and ‘ProtHMSO_M2’respectively represent the use of random single-point mutation and the use of ProtHMSO to mutate two sites, and so on.

### Accelerating genetic algorithm convergence via GA-HMSO

To evaluate the efficacy of integrating ProtHMSO as a component into GA framework, we conducted comparative analyses between conventional GA and two GA-HMSO variants: a fixed crossover mutation rate (GA-HMSO fixed), and a dynamic crossover mutation rate (GA-HMSO dynamic). GA-HMSO fixed retained the default crossover mutation rate of conventional GA, while GA-HMSO dynamic adopted an adaptive strategy, starting with a higher mutation rate to prioritize exploration and then gradually reducing to a lower rate for intensified exploitation. All algorithms were initialized with identical populations derived from three cases in AMP datasets.

As shown in **Fig 7**, conventional GA successfully enhanced the functionality of Case2 and Case3, but failed to improve the activity of Case1 and Case3. In contrast, both GA-HMSO fixed and GA-HMSO dynamic demonstrated consistent superiority across all three cases compared to conventional GA, particularly evident in Case 3, which contains high challenge candidates with low *P*_AMP_ and *P*_MIC_. Notably, as shown in **Table 1**, GA-HMSO dynamic achieved the highest performance with average *P*_MIC_ of 0.82 and 0.83 for Case1 and Case3, outperforming GA-HMSO fixed with average *P*_MIC_ of 0.43 and 0.29 for Case1 and Case3. Besides, GA-HMSO dynamic achieved all average pLDDT values higher than 80 and all average perplexity scores lower than 6 across three AMP cases, which were about 10% better than conventional GA and GA-HMSO-fixed. These results illustrated that by implementing an adaptive mutation schedule, GA-HMSO dynamic effectively balanced global search breadth and local fitness landscape refinement, providing particularly advantageous for navigating the complex, high-dimensional fitness landscape of multi-site protein sequence optimization.

**Fig 7.**
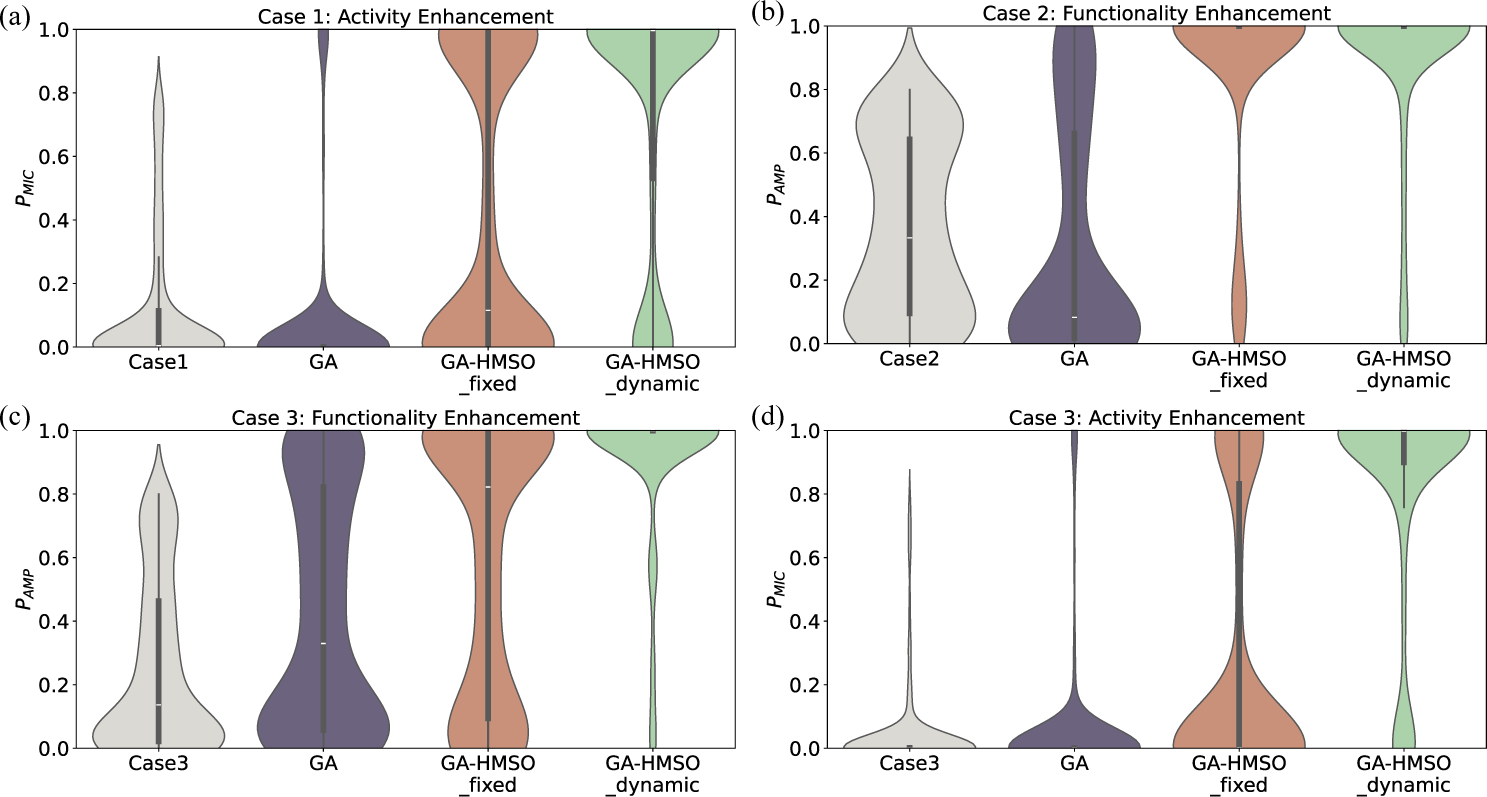
Comparison of functionality or activity enhancement of three AMP cases via GA and two GA-HMSO variants (GA-HMSO fixed and GA-HMSO dynamic). (a) The activity-enhancement of AMPs in Case1. (b) The functionality-enhancement of AMPs in Case2. (c-d) The functionality-enhancement (c) and activity-enhancement (d) of AMPs in Case3.

The performance differentials underscore the critical role of ProtHMSO operators derived from protein language models. By leveraging evolutionary principles alongside learned biological constraints, the hybrid GA-HMSO framework enables efficient exploration of combinatorial sequence space while preserving biological plausibility. This addresses fundamental limitations of conventional evolutionary strategies in computational protein design, particularly in scenarios requiring simultaneous optimization of multiple distant residues.

### Enhancing MCTS exploration via MCTS-HMSO

To evaluate the efficacy of integrating ProtHMSO as a component in MCTS framework, we conducted a comparison between conventional MCTS and its heuristic variant (MCTS-HMSO) across three AMP cases. The exploration efficiency of both methods was systematically analyzed by assessing their optimization outcomes at iterative checkpoints (100, 300, and 500 iterations), as shown in **Fig 8**.

**Fig 8.**
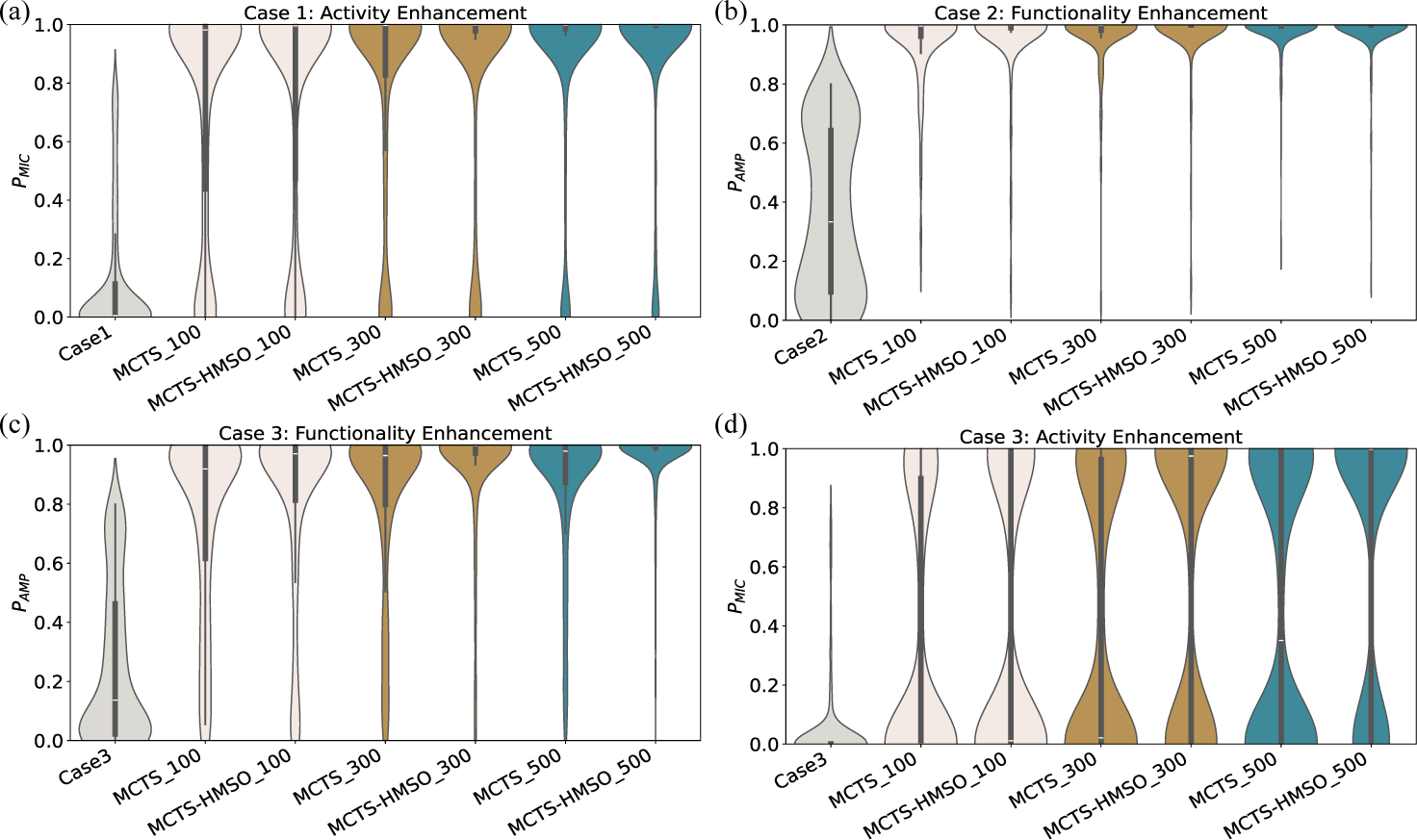
Comparison of functionality or activity enhancement of three AMP cases via MCTS and MCTS-HMSO. (a) The activity-enhancement of AMPs in Case1. (b) The functionality-enhancement of AMPs in Case2. (c-d) The functionality-enhancement (c) and activity-enhancement (d) of AMPs in Case3. ‘MCTS_100’ means the sequence generated by MCTS at 100th iteration, and so on.

Overall, both MCTS and MCTS-HMSO exhibited significant improvement in antimicrobial functionality and activity properties relative to original sequences, and the improvement was further enhanced as the iteration progressed. Moreover, although MCTS has achieved quite high performance, MCTS-HMSO consistently performed better than the MCTS at all three iterative checkpoints across three AMP cases. This superior performance underscores the enhanced exploration efficiency afforded by the HMSO operator, which leverages structural and evolutionary priors from the masked ProtLMs (ESM-2) to guide combinatorial sequence space navigation. By prioritizing beneficial mutations through these learned priors, MCTS-HMSO achieves deeper exploration of the search tree with the same computational budget (i.e. iteration count), thereby enhancing exploration toward high-fitness regions.

Besides, as shown in **Table 1**, MCTS-HMSO also demonstrated consistent improvements in structural stability (pLDDT) and sequence plausibility (perplexity) compared to conventional MCTS. However, the magnitude of improvement was less pronounced than that observed in GA-HMSO. This discrepancy may arise from the inherent constraint of MCTS, which introduces only one mutation per iteration. Such a restriction limits the parallel multi-site exploration of combinatorial mutational spaces, thereby reducing convergence efficiency relative to genetic algorithm-based approaches that permit multi-site modifications in a single iteration.

## Conclusion

In this study, we proposed ProtHMSO, a heuristic multi-site optimization framework for protein sequence design guided by masked protein language models. ProtHMSO enables context-aware directed exploration and was further integrated into genetic and Monte Carlo tree search algorithms (GA-HMSO and MCTS-HMSO) to enhance evolutionary optimization. Experiments on AMP and ProteinGym datasets demonstrate that ProtHMSO consistently improves AMP functionality and activity, effectively explores beneficial mutations in long protein sequences, and accelerates convergence in heuristic frameworks. By coupling evolutionary search with language model–informed mutation guidance, ProtHMSO offers a robust and generalizable strategy for protein engineering while mitigating destabilizing and deleterious substitutions. Future work will explore scalability to larger proteins and integration with physics-based energy functions.

## Acknowledgments

This work is supported by the National Natural Science Foundation of China (62102118), Guangdong Basic and Applied Basic Research Foundation (2025A1515010185), the Shenzhen Colleges and Universities Stable Support Program (GXWD20220811170504001), Shenzhen Science and Technology Program (JCYJ20230807094318038, KQTD20240729102154066).

